# A Simple and Cost-Effective Protocol for Enriching ’*Ca*. Saccharimonadia’ from Human Saliva to Enable Efficient Genome Sequencing

**DOI:** 10.1101/2025.07.17.665291

**Authors:** Clara Bonaiti, Simona Panelli, Giorgia Bettoni, Lodovico Sterzi, Diego Marco Minore, Stella Papaleo, Enza D’Auria, Gianvincenzo Zuccotti, Francesco Comandatore

## Abstract

**Background:** *“Candidatus* Saccharimonadia”, a lineage within the Candidate Phyla Radiation (CPR), is characterized by small cell size, reduced genomes, and an epibiotic lifestyle dependent on bacterial hosts. Despite their presence in diverse environments, including the human oral microbiota, their role in human health remains largely unexplored. Observational studies have linked increased “*Ca*. Saccharimonadia” abundance in human saliva with inflammatory conditions such as periodontitis and inflammatory bowel disease, while recent *in vivo* evidence suggests potential anti-inflammatory properties. Investigating “*Ca*.

Saccharimonadia” is challenging due to their fastidious nature, making standard culturing methods impractical. Current genomic studies rely on either co-cultivation with bacterial hosts or shotgun metagenomics, both requiring advanced technical expertise and costly resources. To address this limitation, we developed an affordable and efficient protocol for enriching “*Ca*. Saccharimonadia” from human saliva samples, facilitating genome sequencing. Our protocol, termed the “*Ca*. Saccharimonadia” Enrichment (SE) protocol, consists of two mandatory phases (detachment from host bacteria and weight/size-based separation) along with an optional DNA degradation phase. We validated this method on two human saliva samples.

**Results:** After SE protocol, the two saliva showed a substantial increase in “*Ca*. Saccharimonadia” DNA concentration. The enriched samples enabled high-quality metagenome-assembled genomes (MAGs) to be obtained *via* shotgun metagenomics.

**Conclusions:** This study presents a cost-effective and scalable approach to studying “*Ca*. Saccharimonadia”, overcoming prior limitations and expanding opportunities for future research on their role in human health and disease.

## Background

*“Candidatus* Saccharimonadia” (formerly known as TM7 or Saccharibacteria) is one of the most studied lineages within the Candidate Phyla Radiation (CPR), a largely unexplored bacterial group which is estimated to encompass approximately 25% of all bacterial genetic variability [1]. “*Ca*. Saccharimonadia” are characterised by[1] small cell size and reduced genomes, which lack several metabolic pathways [2]. Electron microscopy studies have evidenced that “*Ca*. Saccharimonadia” have an epibiotic lifestyle, living on the surface of host *Actinobacteriota [3, 4]*. This suggests a strict metabolic relationship between “*Ca*. Saccharimonadia” and the bacterial host.

“*Ca*. Saccharimonadia” inhabit diverse environments, including water, soil, animals, and are also a recognized component of the human microbiota, residing in the oral cavity, skin, and the gastrointestinal tract [5]. Various studies have shown that “*Ca*. Saccharimonadia” abundance in the oral cavity increases in dysbiotic microbiomes associated with various human inflammatory diseases, including periodontitis, vaginosis and inflammatory bowel diseases [6–8].

On the other hand, experimental evidence revealed that “*Ca*. Saccharimonadia” exhibit anti-inflammatory activities [9], also by inhibiting Tumour Necrosis Factor (TNF)-α production in human macrophages [3]. Coherently, D’Auria and colleagues (2023) found that “*Ca*. Saccharimonadia” are enriched in the oral microbiota of children suffering from food allergy, in comparison to healthy controls [10].

Unfortunately, “*Ca*. Saccharimonadia”, as the other CPR bacteria, are fastidious to culture in standard laboratory conditions [11]. At the state of the art, only a few successful cultivations are reported in literature [3, 12, 13]. This strongly hampers the investigation of “*Ca*. Saccharimonadia” physiology and its impact on human health.

Comparative genomics can aid in reconstructing the metabolic pathways of “*Ca*. Saccharimonadia” and provide insights into their evolution. Currently, genome sequences of these bacteria can be obtained through two approaches: either by co-culturing “*Ca*. Saccharimonadia” with their *Actinobacteriota* host or by using shotgun metagenomics [3, 12, 14]. Both approaches require advanced skills and expensive equipment. The co-cultivation of “*Ca*. Saccharimonadia” can be performed only in very specific conditions, very challenging to achieve. Furthermore, considering that “*Ca*. Saccharimonadia” represents ∼5% in the human oral microbiota [5], the sequencing of their entire genomes by shotgun metagenomics can require a remarkable sequencing depth, increasing the cost and the informatic effort.

At the state of the art, protocols relying on vortexing, filtering and concentrating CPR cells with a final ultra-centrifugation, have been used as preparatory steps for the co-culturing of “*Ca*. Saccharimonadia” with their hosts [12, 13]. However, these approaches have never been used as a culture-independent way to obtain “Ca. Saccharimonadia” genomes.

Building on these protocols, we developed an affordable and efficient protocol for the enrichment of “*Ca*. Saccharimonadia” from human saliva, for genome sequencing purposes. We tested the protocol on two human saliva samples, resulting in a substantial enrichment of the “*Ca*. Saccharimonadia” and the generation of two high-quality genome assemblies, without requiring deep metagenomics sequencing.

## Methods

### Ethics statement

The two saliva samples used in this study were obtained from two children recruited as part of the control population in a study aimed at describing the alteration of oral microbiota in children suffering from food allergies [10]. Participants to the study were recruited at the Pediatric Clinic of the Vittore Buzzi Children’s Hospital (Milan, Italy). The study was conducted according to the Declaration of Helsinki, and all the methods followed the relevant guidelines and regulations. The Ethical Committee of ASST-Fatebenefratelli-Sacco approved the study (Ref. 2021/ST/041). The privacy rights of subjects have been observed. The parents of the recruited children signed an informed consent.

### Human saliva sampling and collection

For each child, two ml of saliva were collected in a sterile container upon awakening, prior to any drinking, eating or teeth brushing. The fresh saliva samples conferred to the hospital were immediately subjected to the *“Ca*. Saccharimonadia*”* enrichment protocol (from here “SE protocol”) described below.

### *“Ca. Saccharimonadia”* enrichment protocol

To enrich “*Ca*. Saccharimonadia” from human saliva samples, we leverage two distinctive traits of these bacteria: the ability to survive on the surface of other bacteria and the small size, along with a resilient cell wall that withstands chemical and physical stresses [15]. The SE protocol is composed of three phases: two mandatory (Phase I and Phase II) and one optional (Phase III). Phase I: detachment of *“Ca*. Saccharimonadia*”* from the surface of host bacteria; Phase II: weight and size-based separation of cells by centrifugation and filtering; Phase III (optional): final removal of free DNA. All these steps are performed manually, prior to DNA extraction.

Phase I (the detachment of “*Ca*. Saccharimonadia”) consists in vortexing one ml of fresh saliva in a two-ml tube for 10 minutes. Phase II (separation of the cells on the basis of weight and size) consists in three main steps: 1) Centrifuge at 2600 x g for 15 minutes; 2) Discard the pellet and filter the supernatant through a 0.45 μm filter; 3) Centrifuge the filtrate at 16,900 x g for 1 hour at 4^°^ C.

These steps are expected to lead to the lysis of several cells, in particular those having a less resistant wall, including eukaryotic cells and gram negative bacteria. To eliminate free DNA, we added an optional third phase using Propidium Monoazide (PMA), a DNA intercalator that degrades DNA upon exposure to 465–475 nm light [16].

Phase III is composed by three steps: 1) the pellet obtained from the centrifugation in Phase II-step 3 is resuspended in 150µl of sterile water by pipetting and vortexing, and left at room temperature for 5 minutes (this step is expected to osmotically lyse mammalian cells); 2) PMA is added to a final concentration of 10µM (7.5µl of the 0.2mM stock) and samples are briefly vortexed, then incubated in the dark at room temperature for 5 minutes; 3) samples are finally placed horizontally on ice, 20cm away from a standard bench top fluorescent light (we used the WESTLITE Flashlight LED Blue 470 nm) for 25 minutes, with rotation every 5 minutes.

The two samples included in the study underwent both a protocol which included only Phase 1 and Phase 2 (from here “SE-short protocol”) and a “SE-long protocol”, which included all three phases. Each SE experiment was performed on two technical replicates for each of the two subjects, and the samples were subsequently stored at -20^°^C.

A graphical representation of the protocols are reported in Figure 1.

**Figure 1.**
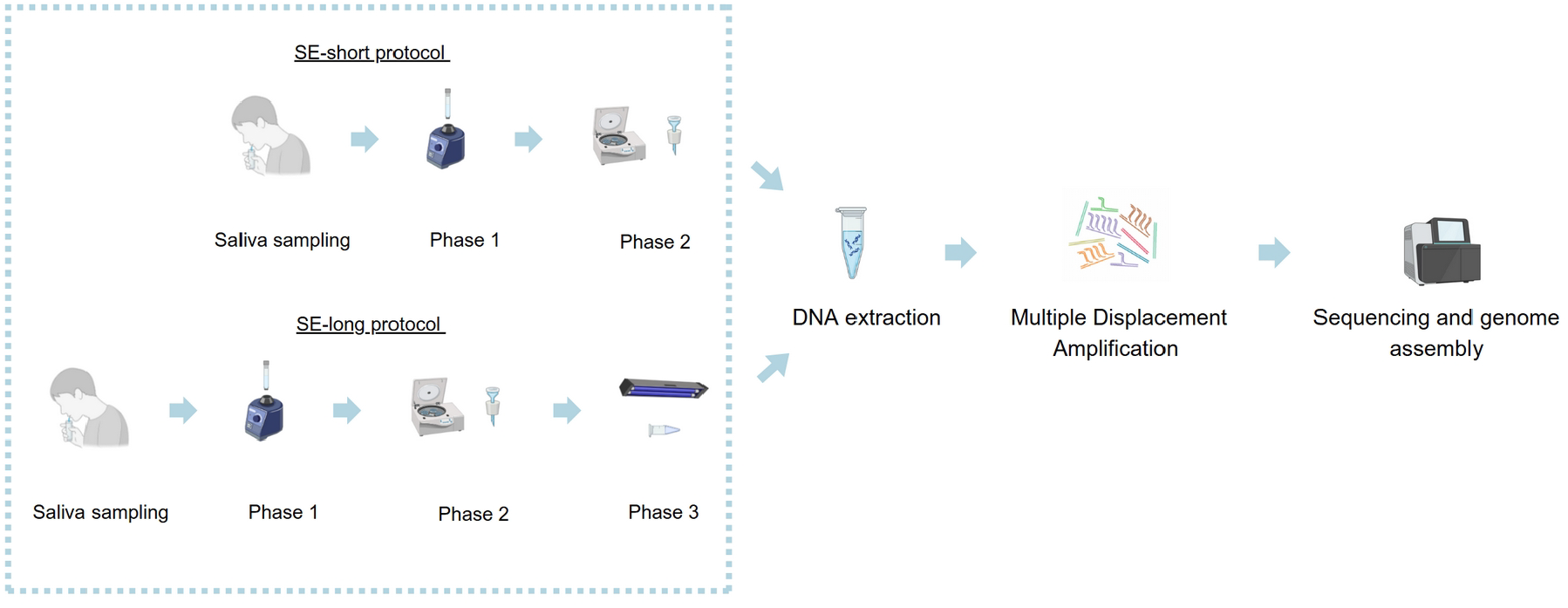
The “Ca. Saccharimonadia” Enrichment (SE) protocol. Graphical representation of the “Ca. Saccharimonadia” Enrichment (SE) protocol, which consists of three phases following fresh saliva sampling: Phase I involves vortexing saliva to detach the target cells; Phase II separates cells by centrifugation and filtration based on size and weight; Phase III (optional) uses Propidium Monoazide (PMA) to eliminate free DNA through light exposure (465–475 nm). The SE-long protocol contains all the three phases, while SE-short only the first two phases. Both SE-short and SE-long protocols are followed by DNA extracted, Multiple Displacement Amplification (MDA), Next Generation Sequencing and genome assembly.

### DNA extraction, quantification and normalization

DNA extractions were performed on original samples (i.e., saliva samples not subjected to the SE protocols), as well as on the samples after SE-short and SE-long protocols. DNA was extracted using the MasterPure™ Gram Positive DNA Purification Kit, following the manufacturer’s instructions, except for the Proteinase K incubation, which lasted one hour instead of 15 minutes. The lysozyme incubation lasted 30 minutes. DNA concentration was assayed with Qubit Fluorometer (Invitrogen, Waltham, Massachusetts, USA). “*Ca*. Saccharimonadia” enrichments obtained by the SE-short and SE-long protocols were then evaluated by V3-V4 16S rRNA sequencing.

### “*Ca*. Saccharimonadia” quantification by V3-V4 16S rRNA amplicon sequencing

Amplicon production was performed as previously described [17]. Libraries were subjected to paired-end sequencing (2 × 300 bp format) on an Illumina MiSeq platform at BMR Genomics (Padova, Italy). The bioinformatics analysis of microbiota sequencing data was based on the Mothur pipeline [18]. Raw FASTQ files were quality-filtered using Trimmomatic [19] and high-quality reads were analysed following the SOP Mothur procedure [20].

### Shotgun metagenomics

One of the primary objectives of the SE protocols is to facilitate the sequencing of “*Ca*. Saccharimonadia” genomes by significantly enriching them in the sample. To evaluate the efficiency of SE protocols in achieving this aim, we performed shotgun metagenomics on the most enriched “*Ca*. Saccharimonadia” sample for each individual included in the study, to assemble the relative “*Ca*. Saccharimonadia” strain genome.

As expected, both the SE-short and SE-long protocols resulted in a substantial reduction of DNA concentration. To mitigate this issue, the DNA extracted after SE protocols was amplified using Multiple Displacement Amplification (MDA) with the Repli-g Single Cell Kit (Qiagen, Hilden, Germany), following the manufacturer’s instructions. The amplified DNA was then sequenced on an Illumina NovaSeq platform (2 × 150 bp paired-end) by Novogene (Munich, Germany).

The low quality reads were trimmed out using the Trimmomatic software [19] and Metagenome-Assembled Genomes (MAGs) were obtained using the MetaWRAP pipeline [21]. The contamination and completeness of the obtained MAGs were then evaluated using the CheckM tool [22]. Then, MAGs were taxonomically assigned using the Forestax tool [23]. The gene content of the MAGs was then compared with a large collection of genome assemblies of the same genus from the Genome Taxonomy Database (GTDB) [24]. The comparison was carried out by orthologues analysis performed using the OrhoFinder tool [25]. Lastly, the metabolic capabilities of the MAGs were investigated by Clusters of Orthologous Groups (COG)-based annotation [26].

To determine the minimum number of reads required to obtain high quality “Ca. Saccharimonadia” MAGs, we repeated the MetaWRAP and Forestax pipelines on four random subsets of 5, 10, 20 and 30 million reads. Once the *Saccharimonadia* MAGs were obtained, the completeness, contamination, genome coverage, contig number, total length and normalized N50 (i.e. N50/total length) were compared.

## Results

### Enrichment of “Ca. Saccharimonadia” from human saliva

The saliva samples collected from the two individuals were subjected to the SE-short and SE-long protocols as described above. DNA from the original samples and after the two protocols were then extracted and subjected to V3-V4 16S rRNA amplicon sequencing to quantify “*Ca*. Saccharimonadia” in the samples (Table 1).

**Table 1.**
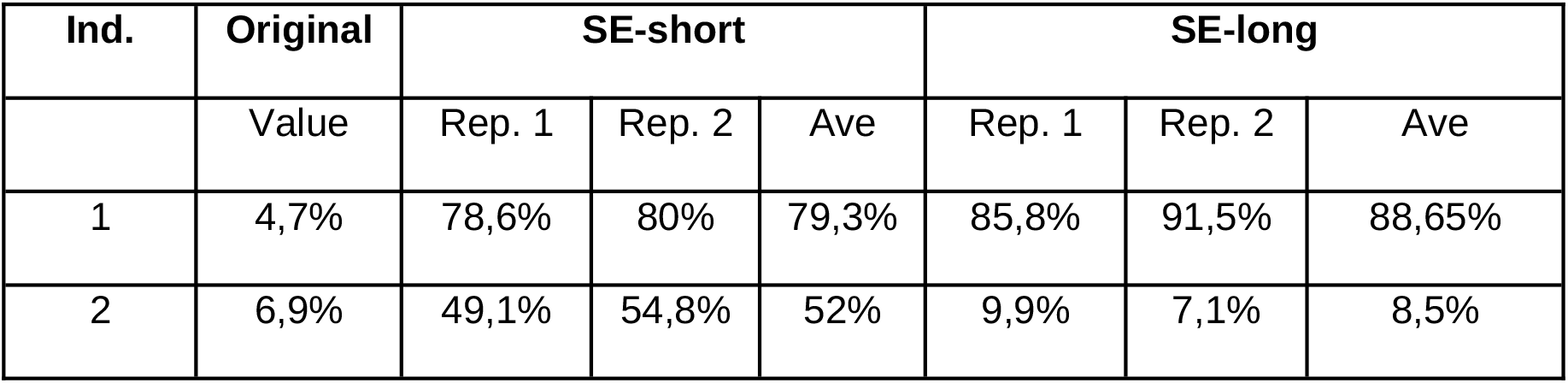
Percentage of “*Ca*. Saccharimonadia” in the samples determined *via* V3-V4 16S rRNA amplicon sequencing The “Original” column refers to “*Ca*. Saccharimonadia” percentage in the collected human saliva samples, while “SE-short” and “SE-long” columns refers to the values obtained after the SE-short and SE-long protocols, respectively. “Rep. 1” and “Rep. 2” refer to the technical replicates.

| Ind. | Original | SE-short |  |  | SE-long |  |  |
| --- | --- | --- | --- | --- | --- | --- | --- |
|  | Value | Rep. 1 | Rep. 2 | Ave | Rep. 1 | Rep. 2 | Ave |
| 1 | 4,7% | 78,6% | 80% | 79,3% | 85,8% | 91,5% | 88,65% |
| 2 | 6,9% | 49,1% | 54,8% | 52% | 9,9% | 7,1% | 8,5% |

### Metagenome-assembled genomes (MAGs)

As reported in Table 1, the samples showing the highest “*Ca*. Saccharimonadia” enrichment were the replicate 2 of the SE-long of the individual 1 (hereafter ‘sample-ind1’) and the replicate 2 of the SE-short of the individual 2 (hereafter ‘sample-ind2’). These two samples were selected for metagenomics.

A total of 53,085,080 and 38,116,642 paired-end reads were obtained for sample-ind1 and sample-ind2, respectively. In the initial stage of the MetaWRAP pipeline, reads were mapped to the human reference genome assembly (Genome Reference Consortium Human Build 38, hg38, GCA_000001405) to distinguish human-derived sequences from those of other organisms, including “*Ca*. Saccharimonadia”. This analysis identified 52,720,686 non-human reads for sample-ind1 (99.3% of the total) and 34,645,562 for sample-ind2 (90.8% of the total). Then, MetaWRAP pipeline subjected the non-human reads to metagenome assembly and Metagenome-Assembled Genomes (MAGs) binning, returning a total of five bins for sample-ind1 and one bin for sample-ind2.

For sample-ind1, two out of five bins resulted to have completeness higher than 70% and contamination below 3%: one with completeness 95.35% and contamination 2.3%, the other with completeness 100% and contamination 2.3%. The taxonomic profiling of the two bins assigned the first one to the genus *Anaeroplasma* (class *Bacilli)* and the second one to the genus *Nanosynbacter* (class *Saccharimonadia*). For sample-ind2, the only bin obtained resulted in the genus *Nanosyncoccus* (class *Saccharimonadia*), with 97.67% of completeness and 0% of contamination.

The gene content and quality of the *Nanosynbacter* genome assembly (MAG sample-ind1) was further assessed by comparing it with 36 Nanosynbacter genome assemblies from the GTDB database (Supplementary Table 1). The same was done for the *Nanosyncoccus* genome assembly (MAG sample-ind2), comparing it with 137 assemblies of the same genus present in GTDB (Supplementary Table 1). The results are reported in Supplementary Table 2 and Supplementary Table 3. For the *Nanosynbacter* genome assembly (MAG sample-ind1) (Supplementary Table 2), most parameters show values around the median of the other assemblies of the same genus in GTDB. About the *Nanosyncoccus* genome assembly (MAG sample-ind2) (Supplementary Table 3), the parameter values are comparable to the highest quality genome assemblies of this genus available in the GTDB database.

### Membrane/Wall proteins annotation

The Clusters of Orthologous Genes (COG)-based annotation of the *Nanosynbacter* and *Nanosyncoccus* MAGs showed differences in the gene content related to bacterial cell wall biosynthesis (Supplementary Table 4). Specifically, the *Nanosynbacter* MAG (sample_ind1) lacked six genes that were present in the *Nanosyncoccus* MAG (sample_ind2), whereas the latter lacked 11 genes found in the former.

### Metagenome-assembled genomes (MAGs) quality and number of reads

For each of the two samples (sample-ind1 and sample-ind2), the MAGs binning and taxonomic profiling were repeated on a random subset of 5 million, 10 million, 20 million and 30 million reads and MAG qualities were compared. The results are shown in Figure 2.

**Figure 2.**
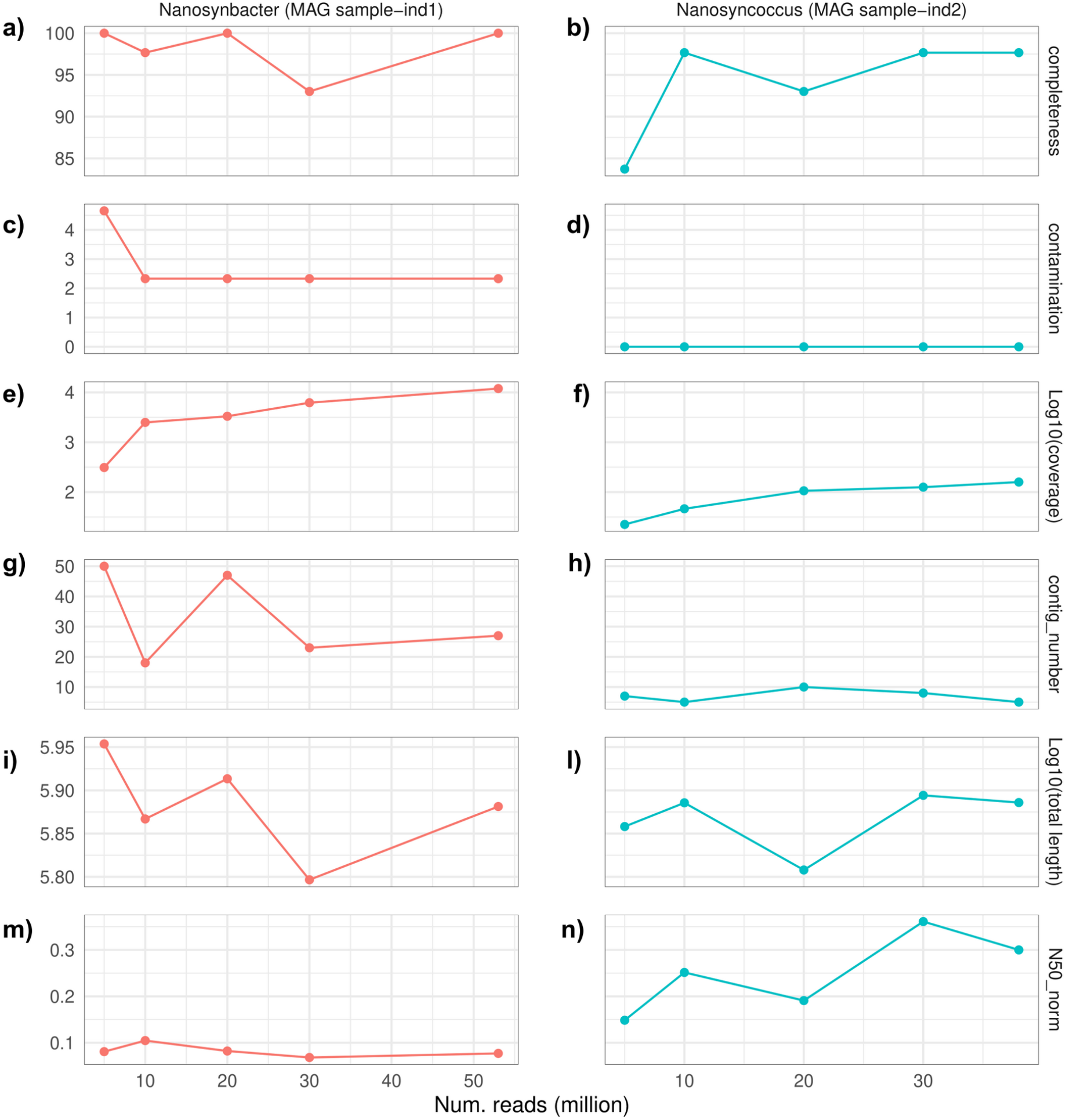
Statistics of MAGs obtained with random subset of metagenomics reads. The plots report completeness (a and b), contamination (c and d), log10 of coverage (e and f), number of contigs (g and h), log10 of MAG total length (i and l), N50 normalized (N50/MAG total length) (m and m), for MAGs obtained by MetaWRAP pipeline using a random subset of 5/10/20/30 millions reads. For each plot, the last value on the right refers to all the sequenced reads (i.e. ∼53 million for Nanosymbacter MAG sample-ind1 and ∼38 million for Nanosyncoccus MAG sample-ind2). The plots a, c, e, g, i, m refer to the MAG obtained from the saliva of the individual 1, after the “Ca. Saccharimonadia” Enrichment (SE) -long protocol. Plots b, d, f, h, i and n, refer to the MAG of the individual 2, even after saliva was subjected to the SE-short protocol.

The *Nanosynbacter* MAG (sample-ind1) reached high completeness and low contamination until 10 million reads. This is also the number of reads that gave the minimum number of contigs, without a strong reduction in the MAG total length and maintaining the coverage above 1,000X. For the *Nanosyncoccus* MAG (sample-ind2), a minimum of 10 million reads were sufficient to obtain high completeness (>95%), ∼50X genome coverage and an assembly total length comparable to the other assemblies obtained with more reads.

## Discussion

Candidate Phyla Radiation (CPR) is a very large but poorly investigated bacterial lineage, found in several ecological niches, ranging from soil to the human body [15].

Observational and experimental studies suggest that CPR bacteria possess immuno-modulating properties [3, 9, 27]. Additionally, D’Auria et al. (2023) found CPR bacteria enriched in allergic children, indicating a potential role in the development of this immune-mediated pathology [10].

The most studied CPR class is “*Ca*. Saccharimonadia”, which is commonly part of the human oral microbiota, even if with frequency ∼5% [5]. This low abundance makes deep and costly metagenomic sequencing necessary to achieve high-quality genome assemblies.

In this work we present a protocol for the enrichment of “*Ca*. Saccharimonadia” from fresh human saliva: “*Ca*. Saccharimonadia*”* Enrichment” protocol, or SE protocol.

This protocol drew inspiration from existing procedures for enrichment of CPR from fresh saliva samples, which rely on vortexing, filtering and concentrating CPR cells with a final ultra-centrifugation. The difference is that, while in these previous works the enrichment was generally preparatory to co-culturing “*Ca*. Saccharimonadia” with their hosts, in our protocol, it is the first step of a procedure that is completely culture-independent and ends in the genomic sequencing of “*Ca*. Saccharimonadia” directly from saliva samples. Another noteworthy point is that we simplified existing protocols by performing the last centrifugation step of Phase II using a standard benchtop microcentrifuge instead of an ultra-centrifuge.

This point makes the protocol feasible by any laboratory equipped with basic instruments. Another original implementation of our protocol is the step of free DNA degradation through PMA-mediated photolysis. This procedure, originally conceived for achieving the depletion of host DNA to increase the rate of bacterial reads in shotgun metagenomics of saliva samples [16], has been tested by us as a way to further increase the “*Ca*. Saccharimonadia” representation in the fresh saliva sample.

The protocol consists of three main phases (two mandatory and one optional) and we propose two protocol variants, called SE-short (which includes the first two phases only) and SE-long (including the three phases). We validated the protocols on two children’s saliva samples, measuring the “*Ca*. Saccharimonadia” enrichment by V3-V4 16S rRNA amplicon sequencing. Then, we assembled two high quality “*Ca*. Saccharimonadia” MAGs using DNA extracted after the SE protocols. Lastly, we evaluated the minimum number of reads required to obtain good quality “*Ca*. Saccharimonadia” MAGs after the enrichment.

Our findings showed that both SE protocols (SE-short and SE-long) effectively enriched *“Ca*. Saccharimonadia*”* from human saliva samples (Table 1 and Table 2), rising up the percentages from ∼5% to ∼50-80%. However, we found that the two individuals carried two different genera of “*Ca*. Saccharimonadia” (i.e. *Nanosynbacter* and *Nanosyncoccus*), that gave a different enrichment response after SE-short and SE-long protocols.

For the Individual 1 a successfully “*Ca*. Saccharimonadia*”* enrichment was observed after both the SE-short and SE-long protocols, and sequencing revealed the presence of a bacterium belonging to the genus *Nanosynbacter*. In contrast, for the Individual 2, which harboured a “*Ca*. Saccharimonadia” bacterium belonging to the genus *Nanosyncoccus*, the SE-long protocol resulted in significantly lower enrichment compared to SE-short.

Both protocols share the first two phases, during which the “*Ca*. Saccharimonadia” are detached from the host bacterial cells (Phase I) and cells with less resistant walls (i.e. eukaryotic and gram negative cells) are expected to be lysed, and removed by filtration (Phase II). At the end of these two phases, DNA from cells with less resistant walls can however remain and be sequenced. The Phase III of the SE-long protocol removes this free-DNA. At the end of the two phases (for the SE-short), or three phases (for SE-long), the sample is subjected to whole DNA extraction and amplification by MDA.

The observed decrease (from 52% to 8.5%) of the “*Ca*. Saccharimonadia” (genus *Nanosyncoccus*) in sample-ind2 after free-DNA removal (Phase III of the SE-long protocol) suggests that a significant portion of the “*Ca*. Saccharimonadia*”* free-DNA (likely released during cell lysis in Phase II) was eliminated in Phase III. This indicates that the “*Ca*. Saccharimonadia*”* species present in Individual 2’s saliva have a non highly resistant cell wall. Conversely, in sample-ind1, the final step of the SE-long protocol had no significant I mpact on “*Ca*. Saccharimonadia*”* (genus *Nanosynbacter*) representation (rising slightly from ∼79% to 88%), suggesting the absence of free DNA from *Nanosynbacter* after Phase II, possibly due to a resistant cell wall. The metabolic characterization of the “*Ca*. Saccharimonadia*”* MAGs obtained from the two samples is coherent with this hypothesis: indeed, MAGs obtained from the individual 2 (which belong to the *Nanosyncoccus* genus) lack several genes involved in the wall biosynthesis.

The MAG obtained after the SE protocols were both of high quality, exhibiting a high level of completeness (>95%) and low contamination (<3%). Additionally, the assembly statistics were comparable or better than those of the “Ca. *Saccharimonadia*” genome assemblies of the genera *Nanosyncoccus or Nanosynbacter* available in public databases.

The MAGs obtained using a subset of 10 million 150×2 paired-end Illumina reads maintained a high level of quality, suggesting that increasing the number of reads would not have a significant impact on the results. This finding is particularly interesting, as this read depth is comparable to that used for Whole-Genome Sequencing (WGS) of bacterial isolates, one order of magnitude lower than what is typically required for metagenomics.

## Conclusions

The protocol presented in this study enables the successful generation of high-quality “*Ca*. Saccharimonadia*”* genome assemblies without the high economic and computational demands of shotgun metagenomics. Moreover, it requires only standard laboratory equipment commonly available in microbiology and molecular biology labs, making it easily implementable. Lastly, the significantly lower sequencing depth needed after applying this protocol further reduces both the computational resources and expertise required for data analysis.

In conclusion, this study presents an easy-to-use, cost-effective protocol for the enrichment of “*Ca*. Saccharimonadia*”*, designed to facilitate the genomic characterization of this elusive bacterial group.

## List of abbreviations

SE protocol: “Ca. Saccharimonadia” enrichment protocol
PMA: Propidium Monoazide
MDA: Multiple Displacement Amplification
MAG: Metagenome-Assembled Genome
GTDB: Genome Taxonomy Database
COG: Clusters of Orthologous Group

## Declarations

### Ethics approval and consent to participate

Participants to the study were recruited at the Pediatric Clinic of the Vittore Buzzi Children’s Hospital (Milan, Italy). The study was conducted according to the Declaration of Helsinki, and all the methods followed the relevant guidelines and regulations. The Ethical Committee of ASST-Fatebenefratelli-Sacco approved the study (Ref. 2021/ST/041). The privacy rights of subjects have been observed. The parents of the recruited children signed an informed consent.

## Authors’ contributions

Made substantial contributions to conception and design of the study: Simona Panelli and Francesco Comandatore; Performed the experiments: Clara Bonaiti, Simona Panelli, Giorgia Bettoni, Stella Papaleo; Performed data analysis: Lodovico Sterzi, Diego Marco Minore; Drafted the manuscript: Francesco Comandatore, Simona Panelli, Lodovico Sterzi, Enza D’Auria, Gianvincenzo Zuccotti

## Acknowledgements

We acknowledge the support of the APC Central Fund of the University of Milano”. We would like to thank Prof. Raffaele De Francesco for his valuable advice during the experimental design.

